# Identification of potential coagulation pathway abnormalities in SARS-Cov-2 infection; insights from bioinformatics analysis

**DOI:** 10.1101/2020.12.07.414631

**Authors:** Sareh Arjmand, Nazanin Hosseinkhan

## Abstract

Abnormal coagulation parameters have been explored in a significant number of severe COVID-19 patients, linked to poor prognosis and increased risk of organ failure. Here, to uncover the potential abnormalities in coagulation pathways, we analyzed the RNA-seq data (GEO147507) obtained from the treatment of three pulmonary epithelial cell lines with SARS-CoV-2. The significant differentially expressed genes (DEGs) were subjected to Enrichr database for KEGG pathway enrichment analysis and gene ontology (GO) functional annotation. The STRING database was used to generate PPI networks for identified DEGs. We found three upregulated procoagulant genes (SERPINE1, SERPINA5, and SERPINB2) belong to the serine protease inhibitor (serpin) superfamily that inhibit tissue plasminogen activator (t-PA) and urokinase plasminogen activator (u-PA) in the fibrinolysis process. In conclusion, we suggest the fibrinolysis process, especially the blockage of t-PA and u-PA inhibitors, a potential target for more study in treating coagulopathy in severe COVID-19 cases.

## Introduction

The current crisis that emerged due to the rapid national and international spread of the new pathogenic SARS coronavirus (SARS-CoV-2) has posed tremendous challenges to the global public health and economy. The number of positive cases and daily deaths is exhibiting a constant progression. Many scientists from different disciplines collaborate to identify possible interventions to reduce the spreading rate and seek effective therapeutics targets. Knowledge obtained from the SARS-CoV (the cause of the previous SARS epidemic) greatly helped to understand the new coronavirus and accelerated its research.

Many of the common and rare clinical features of COVID-19, such as fever, dry cough, myalgia, fatigue, shortness of breath, diarrhea, Acute Respiratory Distress Syndrome (ARDS), vomiting, liver damage, and septic shock, have been well-considered to categorize and monitor disease progression to improve the clinical outcomes ^1, 2^.

After infection, SARS-COV-2 triggers immune responses that, in turn, results in localized lung inflammation. However, in some patients, uncontrolled cytokine responses with unknown causes lead to hyper-inflammation, described as a cytokine storm, and a considerably higher risk of multi-organ failure and death ^3^. A broad spectrum of anti-cytokine and anti-inflammatory therapeutic strategies have been considered to improve patients’ clinical outcomes and prognosis. These immunomodulators must be used in the right time and balanced manner to maintain a sufficient inflammatory response for virus clearance ^4, 5^. However, the hyper-inflammatory state is not the sole factor responsible for lung injuries and organ failure. Rather, the cross-talk between inflammation and coagulation pathways, through which activation of one system amplifies activation of the other, may lead to substantial tissue damage and organ failure in critically ill patients ^6^. Though some studies are starting to show the importance of associated coagulopathy in the prognosis of COVID-19 patients, few clear knowledge has been provided on the hemostasis dysfunction mechanisms in them. Furthermore, inhibition of the coagulation pathway, as a therapeutic target, received less attention ^7^.

Blood coagulation is a dynamic process, triggered as a cascade of proteolytic reaction in which each proenzyme is converted to its active form by an upstream-activated coagulation factor. In the central step of coagulation, Factor Xa converts prothrombin (Factor II) into the active thrombin (Factor IIa). Thrombin works as a master protease that converts the soluble fibrinogen (Factor I) to fibrin (Factor Ia) to form the clots. Thrombin also converts Factor XIII to its activated form (Factor XIIIa), which renders the fibrin monomers to stable cross-linked networks ^8, 9^.

Following clot formation, a separate process called fibrinolysis occurs to degrade the fibrin, restore blood vessel patency, and prevent possible thromboembolism. Plasmin is the main proteolytic enzyme that breakdown the fibrin mesh, leading to the production of circulation fragments (fibrin degradation products (FDP) and D-dimer) cleared by other proteases or by the kidney and liver. Plasminogen is the inactive precursor of plasmin that circulates in free form and is activated, after binding to fibrin, by t-PA and u-PA ^10^. Impaired fibrinolysis may result in a delayed breakdown of clots and predisposes to thromboembolism ^11^. The scheme of coagulation and fibrinolysis is shown in Figure 1.

**Figure 1.**
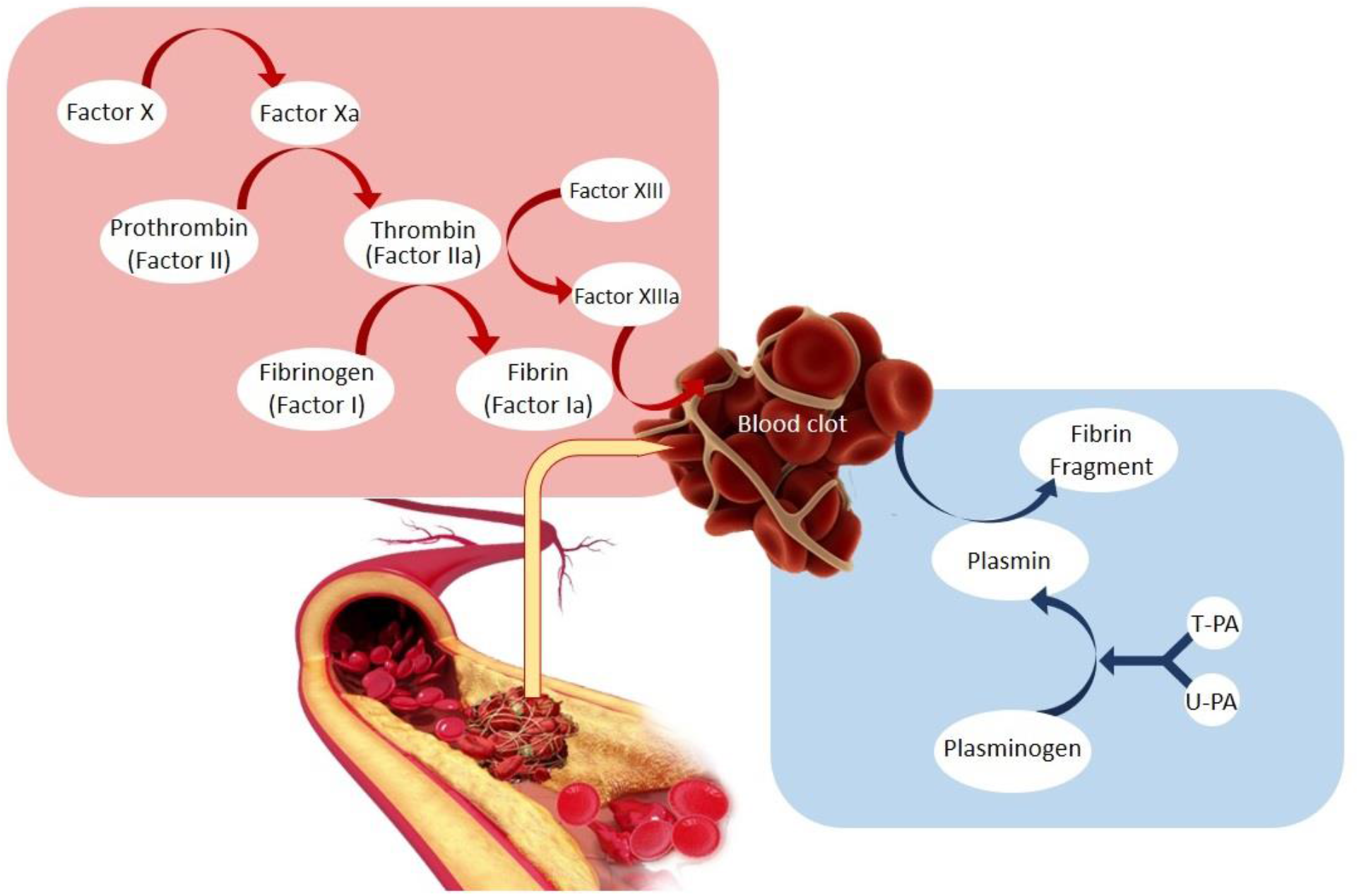
Simplified scheme of blood coagulation and fibrinolysis.

Whole-transcriptome RNA-seq analysis is a high-throughput approach for precise measurement of the expression levels of transcripts ^12^. This method can give an objective view of the molecular mechanism underlying the virus pathogenesis and interaction between host and virus.

The present study aimed to analyze the RNA-seq data (GEO147507) that was provided by Tenoever et al. from primary human lung epithelium (NHBE), transformed lung alveolar (A549) cells, and transformed lung-derived Calu-3 cells infected with SARS-Cov-2 (USA-WA1/2020). DEGs were identified in infected cells 24h post-infection. We compared outcomes with the obtained results from the infection of cells with Avian Influenza Virus (AVI) subtype H1N1, the bird flu with pandemic potential for humans. We analyzed the dataset to uncover the potential abnormalities in coagulation pathways that occur due to the infection with SARS-CoV-2. We hope that this study offers new insights into the potential mechanisms involved in SARS-CoV-2 pathogenesis and would prove a useful theoretical basis for future biological and clinical applications.

## Materials and methods

### Acquisition of microarray data

Transcriptional response to SARS-CoV-2 infection dataset GSE147507 submitted by Tenoever BR., and Blanco-Melo D. based on GPL18573 platform (Illumina NextSeq 500) was downloaded from GEO database, in the National Center for Biotechnology Information (NCBI).

### Differential Expression Analysis

FASTQ files were introduced to FASTQC algorithm to check the presence and the type of adaptor contaminations within FASTQ files. Trimming of FASTQ files was then carried out using Trimmomatic algorithm. We used GRCH37 (hg19) version of the reference genome for mapping reads using “*Bowtie for Illumina*” algorithm. Read counts for each feature (gene) were then calculated using “*HTSeq-count*” algorithm. Finally, we employed “*DESeq2*” algorithm on the resulted read counts to get the list of deregulated mRNAs. Applying the adjusted p-value ≤0.05, the most statistically significant up and down-regulated genes were selected. To identify deregulated genes, the biologically significant changes between mock and treated samples, |logFC| >= ±0.58 cutoff was applied, which equals to at least 1.5 folds increase or decrease in expression values.

### GO and KEGG enrichment analysis

The significant DEGs were mapped into Enrichr database for KEGG pathway enrichment analysis and GO functional annotation. A *p* < 0.05 depicts significant enrichment.

### PPI network investigation

The STRING database was used to predict the interaction relationship between DEGs corresponding to coagulation. Only the interactions with experimental support were chosen for PPI data acquisition. The reliability of PPI interactions were checked based on the provided experimental information. The integrated PPI network was visualized and analyzed using cytoscape v3.8.2.

## Results

### Identification of DEGs and enrichment analysis

The list of DEGs in all cell types treated with SARS-CoV-2 and H1N1 is provided in supplementary 1. The identified DEGs enriched in the coagulation pathway are shown in Table 1.

**Table 1.** The list of DEGs related to the coagulation pathway in cells treated with SARS-CoV-2 and H1N1

| Cell line-virus-MOI | Up-regulated (LogFC) | Down-regulated (LogFC) |
| --- | --- | --- |
| A549-SARS-CoV-2-MOI2 | SERPINE1 (4.11) | ---- |
| Calu3-SARS-CoV-2-MOI2 | SERPINA5 (1.63) | ---- |
| NHBE-SARS-CoV-2-MOI2 | SERPINB2 (1.24) | ---- |
| A549-H1N1-MOI5 | PLAUR (1.33);TFPI(0.92) | PROC (-1.31) |
| NHBE-H1N1-MOI5 | SERPING1 (1.28) | ---- |

### PPI network

The identification of proteins that interact directly with DEGs involved in the coagulation pathway could help understand the molecular mechanism of hypercoagulation in COVID-19 patients. In the present study, two PPI networks were built for the results of SARS-CoV-2 (three upregulated genes) and H1N1 (three upregulated and one downregulated genes) separately and compared (Figure 2).

**Figure 2.**
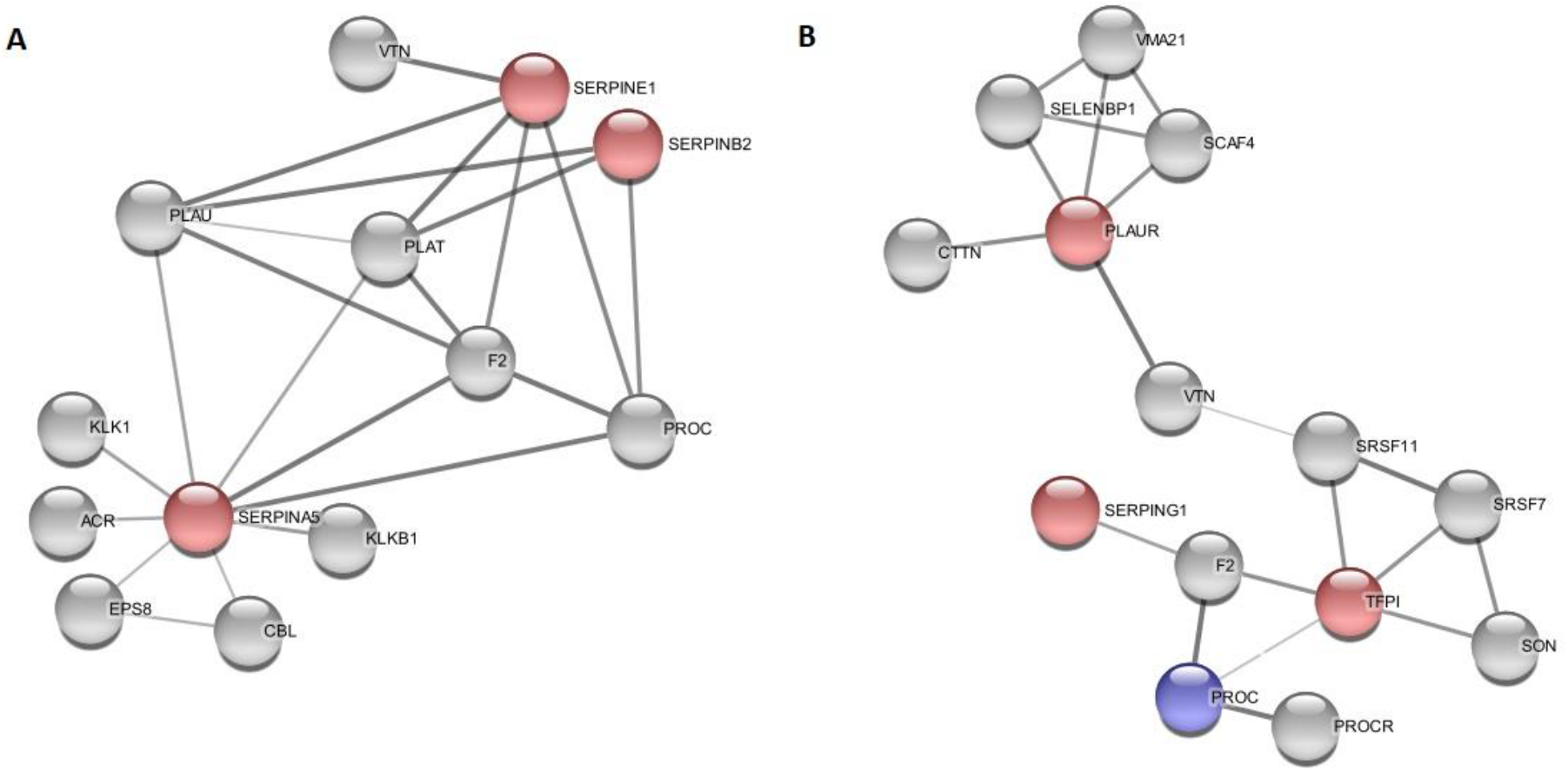
Proteins physically interacting with coagulation associated DEGs in A) SARS-CoV-2 virus, and B) H1N1. PPIs were revealed based on the STRING database with a score =0.4 (median confidence), and nodes represent DEGs in the PPI interaction. The red nodes are related to upregulated and the blue node to the downregulated ones.

## Discussion

Based on the severity of the symptoms, patients with COVID-19 is classified into four levels; mild, moderate, severe, and critical. While mild patients, which constitute a large number of patients, exhibit weak or no symptoms, the critical patients quickly progress to destructive symptoms such as ARDS, coagulation dysfunction, multiple organ failure, and loss of consciousness ^2^.

A significant number of cases with severe COVID-19 infection experience clotting problems that were more likely to receive ICU admission. It is found that blood clots are closely linked to the increased risk of organ failure and death ^4^.

Coagulation is an integral part of innate immune responses that slows down the circulation of invasive microorganisms and limits immune components’ loss. In turn, the natural anticoagulants modulate their function to limit excessive coagulation, maintain the hemostatic balance, and reduce tissue destructions ^13^. Disruption of this well-regulated procoagulant-anticoagulant balance leads to pathologic conditions such as thrombosis or bleeding ^14^.

Accumulated evidence reveals that abnormal coagulation parameters are often seen in severe COVID-19 with a poor prognosis. Lee et al. reported that 20-55% of hospitalized patients with COVID-19 have laboratory evidence of coagulopathy and significantly worse prognosis ^15^. Tang et al. also showed that 71.4% of non-survivors from COVID-19 and only 0.6% of survivors fulfilled the criteria of disseminated intravascular coagulation (DIC) in their hospitalization period ^16^. Cui et al. indicated that the conventional coagulation parameters of COVID-19 ICU patients were significantly higher than those of non-ICU ones and that the incidence of venous thromboembolism (VTE) in patients with COVID-19 was 25% ^17^.

Coagulation dysfunction has been proposed as the complications of COVID-19, which can ultimately lead to ischemic stroke, myocardial injury, and pulmonary thromboembolism, particularly in individuals with underlying comorbid diseases ^18-21^.

The site of thrombosis in the majority of severe COVID-19 is the lungs ^22^. Postmortem examination of lungs in COVID-19 patients has revealed widespread microthrombotic occlusion of small pulmonary vessels ^23, 24^. The incidence of pulmonary embolism has been measured by 23-30% in severe COVID-19 patients ^25^. The pulmonary thromboembolism (PTE) is diagnosed in hospitalized COVID-19 patients by observation of the sudden onset of oxygenation deterioration and shock ^26^. It seems that a variety of potential risk factors, including infection, the immobilization of patients due to prolonged ICU admission, mechanical ventilation, hypoxia, and the hypercoagulable state predisposes them to the high risk of PTE ^27, 28^.

Early detection of life-threatening blood clots in the lung could lead to rapid treatment intervention and reduce the mortality rate. However, a specific diagnosis of PTE is challenging. Understanding the molecular mechanism of SARS-CoV2 behavior and host response to the virus will greatly help develop antiviral therapies and diagnosis modalities. However, there is a paucity of information regarding this issue. Belanco-Melo et al., performed an in-depth analysis of the transcriptional response to SARS-CoV-2 compared with other respiratory viruses in cells and animal model (GEO147507) ^29^. Here we investigated the RNA-seq data obtained from the three respiratory cell lines (A549, NHBE, and Calu-3) and focuses on the expression changes observed in genes related to the coagulation pathway.

In low-MOI infection (MOI,0.2) for the A549 cell line, no DEGs associated with the coagulation pathway was identified. Hence, we used the results of SARS-CoV-2 virus infection with MOI 2 for all three cell lines. KEGG pathway enrichment analysis results indicated that the complement and coagulation cascade in all three cell lines was significantly enriched in SARS-CoV-2 treated cells. In this upregulated pathway, we focused on the genes directly involved in clot formation and degradation. Three genes, SERPINE1, SERPINA5, and SERPINB2 were upregulated in three cell lines, A549, Calu3, and NHBE, respectively. All three identified genes belong to the serpin superfamily of protease inhibitors.

Different serine proteases involve in the coagulation and fibrinolysis pathways that are regulated by feedback loops or serpins. So, improper function of serpins can cause either pathological thrombosis or bleeding ^30^. Serpins represent a broad family of proteins with a wide range of cellular locations and functions. They share a conserved three-dimensional structure that is crucial in their function. Most serpins irreversibly inhibit the target serine proteases, and the stoichiometry value of inhibition is 1 ^31^.

A549 is a pulmonary adenocarcinoma cell line, widely used in lung respiratory and cancer research as a model for lung alveolar epithelium ^32^. Jain et al. indicated that human lung tumor-derived alveolar epithelial cell lines could exhibit pro-thrombotic activities and plays a critical role in pulmonary thrombosis induced by lipopolysaccharide (LPS) endotoxin. So LPS, which is used to induce an experimental inflammation model, indirectly stimulates intravascular thrombosis by activating the alveolar epithelium rather than acting directly on endothelium ^33^. Furthermore, it was reported that alveolar epithelium is capable of modulating intra-alveolar coagulation and may contribute to intra-alveolar fibrin deposition by exposure to inflammatory stimuli ^34^. Hence the results obtained from the treatment of A549 cell line with SARS-CoV-2 could give us useful information about the mechanism of lung clot formation in COVID-19. The result of expression analysis indicated that when A549 is treated with SARS-CoV-2, the most upregulated gene is SERPINE1, with a more than 17 fold increase in expression (logFC=4.1). According to pathway enrichment, SERPINE1 is the only identified gene involved in the coagulation pathway after the treatment of A549 with SARS-CoV-2.

SERPINE1 gene encodes the plasminogen activator inhibitor 1 (PAI-1), which is the most important inhibitor of t-PA and u-PA in the fibrinolysis process. PAI-1 is secreted as an active protein and, by binding to t-PA and u-PA, blocks the activation of plasminogen to plasmin, and therefore the fibrin clot hydrolysis. It was indicated that the production and secretion of PAI-1 could be induced by stimulators like thrombin, endotoxin, cytokines, and chronic inflammation ^35, 36^. This inhibitor is required for the downregulation of fibrinolysis and the control of blood clot degradation ^37^. High levels of circulating PAI-1 was shown to be associated with some thrombotic diseases and cancers ^38, 39^, and transgenic mice overexpressing native human PAI-1 developed spontaneous thrombosis ^40^. Overexpression PAI-1 has been known as a biomarker of aging-associated thrombosis and a variety of other pathologies (e.g., obesity, hyperinsulinemia, type-2 diabetes, coronary heart diseases, a decrease of immune responses, a proliferation of inflammation, and vascular sclerosis/remodeling) ^41, 42^. Based on this information, it seems that this inhibitor could be considered a notable target for inhibition in COVID-19 patients at the risk of developing pulmonary thromboembolism. It is noteworthy that already some fibrinolytic drugs, such as heparin and tPA, are administering to remedy the thrombotic pathology in COVID-19, and it is suggested this therapy is effective in some patients ^22, 43^. Suppressing of PAI-1 (using specific antibodies, antisense oligonucleotides, peptide antagonists, etc.) has been suggested for further studies as a potential therapy for complications such as renal fibrogenesis, breast cancer, and vascular diseases ^39, 41, 44^. Furthermore, a wide range of small molecules with anti-PAI-1 activity have been isolated from natural sources that showed satisfactory results *in vitro* and with some *in vivo* models ^45^. Interestingly, the treatment of A549 with the H1N1 AVI virus (with higher MOI (=5) compared to the SARS-CoV-2) showed completely different results. The analysis indicated that AVI induces the expression of two anticoagulants gene (plasminogen activator urokinase receptor (PLAUR), and tissue factor pathway inhibitor (TFPI)) and downregulation of protein C (PROC), the zymogen of activated protein C (APC).

The results of the analysis on the other bronchial epithelial cell (NHBE) with the same MOI (=2) indicated the ∼2.24 fold (logFC=1.24) overexpression of SERPINB2 that is involved in the coagulation pathway. SERPINB2 or plasminogen activator inhibitor 2 (PAI-2) is a well-described coagulant factor that inactivates u-PA and t-PA, and impaired fibrinolysis. It has been shown that the expression of PAI-2 is associated with placental tissue and monocyte macrophages ^46^. PAI-2 level is normally very low in blood but is drastically elevated during pregnancy, partially explaining the higher risk of thrombosis during pregnancy ^47^. It is also markedly induced by proinflammatory mediators, such as tumor necrosis factor-alpha (TNF-*α*) and LPS, and a variety of cytokines and growth factors that are involved in cell differentiation ^48, 49^. PAI-2 was also detected from the bronchoalveolar lavage of patients with ARDS and different forms of interstitial lung diseases, where the fibrinolysis is hardly detectable. The considerable decrease in fibrinolysis under the mentioned circumstances is due to the overexpression of both PAI-1 and PAI-2 ^50, 51^. To the best of our knowledge, inhibition of PAI-2 (unlike PAI-1) has not yet been studied to improve coagulopathies. Herein, inhibition of the PAI-2, besides PAI-2, is suggested as another exciting target for studies related to reversal fibrinolysis to the normal condition in COVID-19 patients. In NHBE cells, just like the A549, treatment with AVI virus led to contradictory results. AVI induced an overexpression in the SERPING1 or plasma protease C1 inhibitor (C1-INH), which is a protein with a dose-dependent anticoagulant effect in human whole blood ^52^.

Treatment of Calu3, another lung epithelial cell used in this study, with SARS-CoV-2 led to a significant increase (∼1.3 fold, LogFc=1.63) in serpinA5 or plasminogen activator inhibitor 3 (PAI-3), known as protein C inhibitor (PCI) as well. PAI-3 is a multi-functional serpin playing several roles depending on organs, tissues, and species ^53^. This protein affects blood coagulation and fibrinolysis in different ways. Initially, it was found to be the inhibitor of activated protein C (APC). APC is a natural anticoagulant that down-regulates the blood coagulation by proteolytic degradation of FVa and FVIIIa ^54^. PAI-3 also inhibits u-PA, t-PA, and several other serine proteases involved in blood coagulation and fibrinolysis ^55^.

Overall, here we found that SARS-CoV-2 treatment led to the upregulation of three procoagulant genes in pulmonary epithelial cells. Interestingly, the three upregulated genes’ common feature is their involvement in the fibrinolysis process and inhibition of clot degradation. As it is clear from the PPI network, all three detected serpin interact with u-PA (in Figure 2 is shown as PLAU), and two of them, SERPINE1 and SERPINA5, was induced by thrombin. The equal experiment using H1N1 viruses that affects the pulmonary system as well led to contradictory results, and the detected upregulated genes have known for their anticoagulatory effects through the inhibition of the coagulation pathway and inhibition of clot formation. From the result of PPI for H1N1 it seems that this virus does not affect fibrinolysis directly.

In conclusion, we suggest the fibrinolysis process, especially the blockage of t-PA and u-PA inhibitors, a considerable and potential target for preventing and treating coagulopathy in severe COVID-19 cases.

## Notes

### Competing Interest Statement

The authors have declared no competing interest.

